# Cell confinement initiates a delayed but heritable loss of chromosomes

**DOI:** 10.64898/2026.02.03.703566

**Authors:** Steven H. Phan, Mai Wang, Kayla R. Snare, Jonah Kandell, Jeneille S. Deans, Panteleimon Rompolas, Guilherme P.F. Nader, Dennis E. Discher

**Affiliations:** Molecular and Cell Biophysics Lab, University of Pennsylvania, Philadelphia, PA 19104, USA; Department of Pathology and Laboratory Medicine, Children’s Hospital of Philadelphia, Philadelphia, PA, USA; Department of Cell and Developmental Biology, University of Pennsylvania, Philadelphia, PA 19104, USA

**Author notes:** Senior author and Lead contact (Dennis E. Discher).

**Keywords:** Mechanobiology, Compression, Genetics, Aneuploidy, Cancer, Evolution

## Abstract

Heritable genetic changes continually arise in cancer, especially in solid tumors where cells sometimes appear compressed. Rare heritable losses of chromosomes in live cells are quantified here with chromosome reporters (ChReporters) that reveal similar levels of loss after imposing a threshold level of confinement. Compression to ∼60% of interphase height ruptures few nuclei compared to deeper compression but perturbs mitotic spindles and prolongs pro/metaphase. Chromosome mis-segregation into micronuclei is discovered only *after* release from modest confinement, but arrest and death predominate. All such effects are phenocopied by Nocodazole washout that generates a ‘memory’ of prolonged mitosis, and effects differ from the rapid induction of micronuclei by a spindle assembly checkpoint inhibitor and by a clinical CDK4/6-inhibitor of cell cycle entry. Single-cell-RNA-sequencing confirms chromosome loss days after confinement and reveals persistence of chromosome segregation pathways. Chromosome losses as mitotic memories of confinement ultimately address knowledge gaps in mechanobiology and cancer evolution.

## INTRODUCTION

Heritable genetic changes to individual cells initiate cancer, continually arise, and drive evolution including treatment resistance. Matrix-rich, stiff solid tumors intriguingly exhibit the most genetic changes including chromosome losses and gains that alter expression of 100’s to 1000’s of genes,^1^ but whether mechanical factors such as cell confinement by stiff matrix and cell crowding *cause* heritable mutations remains unclear. Some hallmarks of cancer – including proliferation, invasion, epigenetics, and immune dysfunction – certainly exhibit a degree of mechano-sensitivity.^2–10^

Compression of cells in solid tumors can be controllably studied *in vitro* with devices that impose precision confinement.^11,12^ Strong confinement of interphase cells *in vivo* and *in vitro* causes nuclear envelope rupture and DNA damage with possible outcomes being cell arrest, death, and maybe mutation via mis-repair. Mitotic chromatin in stiff solid tumors appears compressed to heights of interphase nuclei whereas mitotic heights of chromatin in 2D culture typically increase relative to interphase (**Fig.S1A**). Confinement *in vitro* causes spindle distortions and aberrant mitosis in stationary and migrating cells.^13–16^ However, migration under confining and constricting conditions will select for subsets of cells,^17^ and given the genetic heterogeneity of cell culture lines,^18^ any selection confounds sequencing methods that try to assess the *induction* of heritable changes such as chromosome losses and gains.^15,19,20^

Here we quantify the kinetics of *de novo* chromosome losses caused by precise confinement using live-cell chromosome reporters (ChReporters). These enable visualization and quantitation of the *first* losses among >10,000 cells per replicate.^19^ Single cell sequencing experiments cannot track the same rare cell in space-time and might track only a couple dozen cells per condition.^21^ We focus on losses because they precede gains in human tumors^22^ and because chromosomes with key tumor suppressor genes are lost at high frequency across cancers.^23^ Use of ChReporters in stiff solid tumors versus 2D cultures in our recent studies^19^ showed greater ChReporter loss *in vivo* (**Fig.S1A**), and given the noted compression of mitosis *in vivo*, we hypothesized that confined mitosis rather than interphase nuclear rupture drives chromosome loss. Regardless, the sensitivity of chromosome loss to compression is unknown.

Chromosome losses and gains are possible outcomes of chromosome mis-segregation into micronuclei,^24,25^ but *heritable* losses or gains are prevented by cell cycle arrest or death that also result from mis-segregation.^5^ Perturbing the spindle or its linkages to chromosomes often generates micronuclei, and disruption of the spindle with Nocodazole (Noc), for example, prolongs mitosis until washout, which subsequently increases mis-segregation and micronuclei through *fatigue* of the cohesion between chromatids.^21,26^ Prolonged mitosis also increases the likelihood of daughter cell arrest/death as one facet of a ‘memory’ for many normal and cancerous lines.^27^ Inhibition of the spindle assembly checkpoint kinase MPS1 accelerates mitosis (rather than prolonging it) but has some similar outcomes as Noc washout.^28^ as does a clinically relevant inhibitor of CDK4/6 that blocks cell cycle entry.^29^ Whether micronuclei form during or after confinement of mitotic cells remains unknown.

We address a significant gap in mechanobiology and cancer evolution by assessing confinement effects on interphase and mitotic cells (**Fig.1Ai**). Drugs noted above probe the kinetics and outcomes. We primarily use a lung cancer ChReporter line for Chr-5, with mono-allelic expression on RFP-LMNB1, but show similar results for a gene on the much smaller Chr-19. ChReporter-5 negative cells lack a key tumor suppressor and eventually select for proliferation,^18^ but our focus here is the initial induction of Chr loss.

## RESULTS

### Confinement of mitotically rounded cells drives heritable Chromosome loss

To assess whether chromosome loss depends on confinement height, we first measured mitotic and interphase chromatin heights of ∼14µm and ∼10µm, respectively, for our A549 ChReporter cells (**Fig.1Aii**). These cells were then confined to heights of 2, 6, 10 or 19µm after sorting for ChReporter-positive cells (<0.01% negative) (**Fig.S1B,C**). After 16h of confinement, which is about half the cell cycle in standard 2D culture, a recovery period of 2 days allows 1-2 more divisions, which dilutes and dims RFP-LMNB1 protein in cells that have lost chromosome-5.^19^ The approach models tumor suppressor loss in that cells lose chromosomal DNA for the gene, but RNA and protein must then also degrade or dilute via divisions. We quantified ChReporter-negative cells in triplicate by flow cytometry on >10,000 cells (**Fig.1B**), and the 2D Ctrl’s typical variation between experiments (0.08% to 0.12%) equates to differences of only ∼1-per-1000 cells from flow cytometry on 10-100k cells. For all confining heights ≤10µm, ChReporter loss was always ∼2-fold higher than 2D Ctrl’s (**Fig.1Bii-iii**). Therefore we focused on the intermediate height of 6µm.

**Figure 1.**
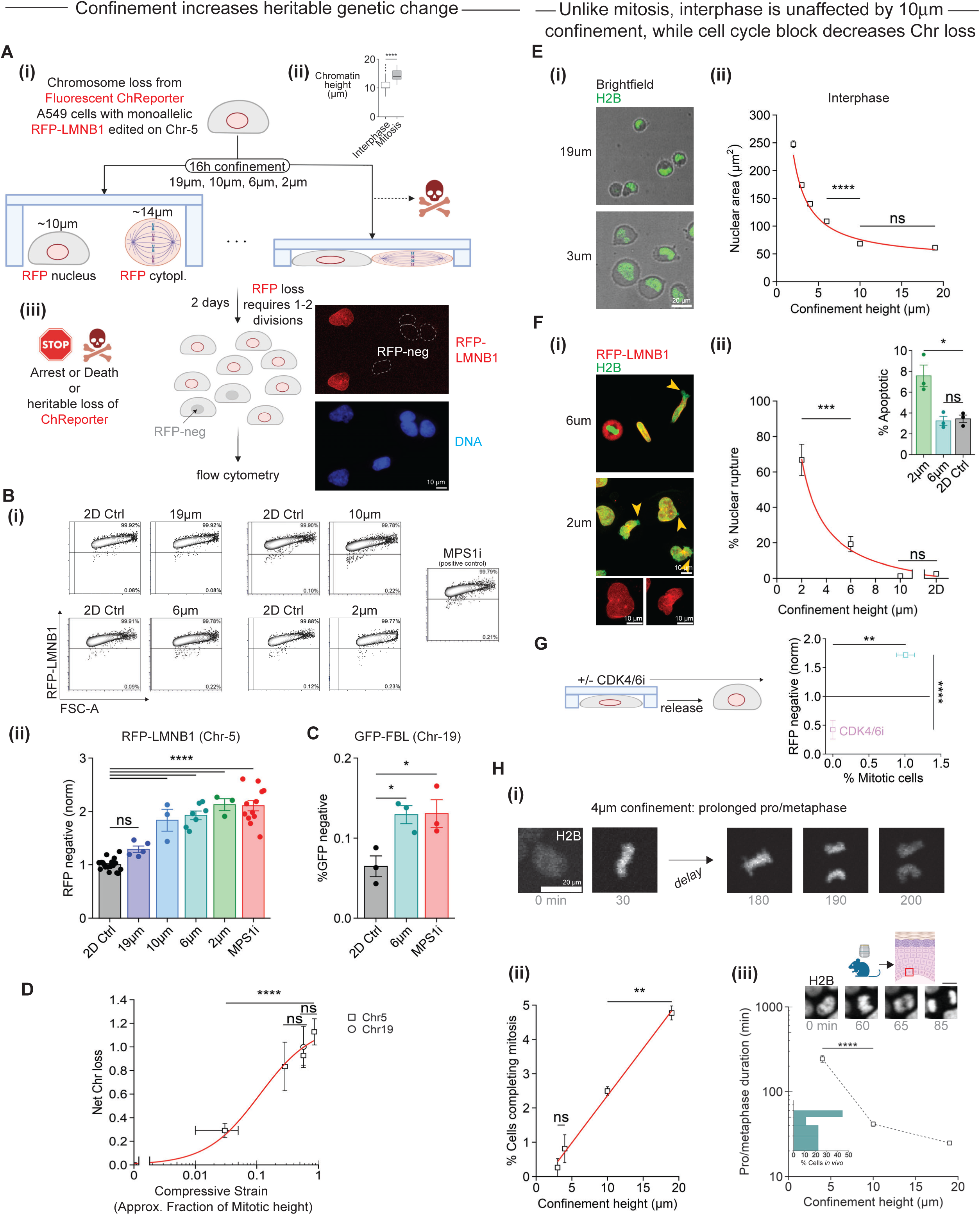
Compression overnight increases heritable genetic changes as well as nuclear spreading and mitotic suppression. Also see Figure S1. **(A)** Experimental design for confinement of mitotic cells or both mitotic and interphase cells to precise heights. **(i)** Freshly sorted A549 Chr-5 ChReporter (RFP-LMNB1) cells were confined for 16h. **(ii)** Chromatin heights differ for interphase and mitotic pro/metaphase. (n≥46 cells; Mean, SEM; ordinary_one-way_ANOVA_and_Tukey’s_ multiple_comparisons_test; ****p<0.0001). **(iii)** Possible fates of death, arrest, and viable colony of ChReporter-negative cells several days after confinement. Scalebar = 10μm. **(B)** Flow cytometry of ChReporter cells after recovery from confinement. **(i)** Representative plots. **(ii)** Quantification of confined ChReporter-negative cells (normalized to 2D control). Cells were also treated with 50nM MPS1i (reversine) to induce chromosome mis-segregation for 16h followed by 2 days recovery (positive control). (n=3 biological replicates; Mean, SEM; ordinary_one-way_ANOVA_and_ Tukey’s_multiple_comparisons test; n.s. not significant, **p<0.01, ***p<0.001, ****p<0.0001). **(C)** Quantification of confined Chr-19 ChReporter (GFP-FBL) negative cells. (n=3 biological replicates; Mean, SEM; ordinary_one-way_ANOVA_and_Tukey’s_multiple_comparisons_test; *p<0.05). **(D)** Two-state fit *y* = *ax* / (*b* + *x*) = *a*[1 + *be*^−ln(*x*)^]^−1^. Baseline-subtracted loss versus ‘Relative strain’: *x* = (1 – confinement height / unperturbed height) using 14µm unperturbed height; assuming 19µm strains large cells to *x* = 0.03+/-0.02. (n≥3; Mean, SEM; ordinary_one-way_ANOVA_and_Tukey’s_multiple_ comparisons_test; n.s. not significant, ****p<0.0001). **(E)** Interphase nuclei spread under confinement. **(i)** Brightfield and GFP-H2B images. Scalebar = 20μm. **(ii)** Nuclear area versus confinement height. (n≥100 cells; Mean, SEM; ordinary_one-way_ANOVA_and_ Tukey’s_multiple_comparisons_test; n.s. not significant, ****p<0.0001). **(F)** Nuclear rupture and death. **(i)** GFP-H2B and RFP-LMNB1 reveal rupture (arrows). RFP-LMNB1 is lacking in DNA blebs. Scalebar = 20μm. **(ii)** Nuclear rupture versus confinement height. (n=3 biological replicates; Mean, SEM; ordinary_one-way_ANOVA_and_Tukey’s_ multiple_comparisons_test; n.s. not significant, ***p<0.001). **(inset)** Analysis of cell death after 16h of confinement. (n=3 biological replicates; Mean, SEM; ordinary_one-way_ANOVA_and_Tukey’s_multiple_comparisons_test; n.s. not significant, *p<0.05). **(G)** CDK4/6i prevents cells from exiting G1 and is sustained both during (16h) and after (48h) confinement. Quantification of ChReporter loss and mitotic cells for +/− CDK4/6i (10µM) and 6µm confinement normalized to 2D control. (n≥3 biological replicates; Mean, SEM; ordinary one-way ANOVA and Tukey’s multiple comparisons test; n.s. not significant, **p<0.01, ****p<0.0001). **(H)** Mitotic suppression by confinement. **(i)** Time series of a cell undergoing protracted mitosis while confined. Scalebar = 20μm. **(ii)** Quantification of mitotic events over 16h confinement. (n≥3 biological replicates; Mean, SEM; ordinary_one-way_ANOVA_and_ Tukey’s_multiple_comparisons_test; n.s. not significant, **p<0.01). **(iii)** Quantification of pro/metaphase to anaphase duration. (19µm: n=30 cells, 10µm: n=27 cells, 4µm: n=16 cells; Mean, SEM; ordinary_one-way_ANOVA_ and_Tukey’s_multiple_comparisons test; n.s. not significant, *p<0.05). **(inset)** Quantification of pro/metaphase duration of skin stem cells *in vivo* (20 cells in n=2 experiments). Scalebar = 10μm.

Cultures treated for 16h with an inhibitor of spindle assembly checkpoint kinase MPS1 served as a positive control and also showed a ∼2-fold increase in ChReporter loss. A similar ∼2-fold increase was observed for a ChReporter made with the smaller Chr-19 (A549 cells with GFP-FBL)^19^ (**Fig.1C,S1D**), and for a different cell line (H23 lung cancer cells with GFP-LMNB1) (**Fig.S1E**). Copy number alterations (CNA) per division were estimated by multiplying our measured chromosome losses (∼0.001; **Fig.1B,C**) by the ∼60-70 chromosomes in A549 cells^18^ which gives ∼0.1 CNA per division. This is close to the 0.2-0.3 CNA per division based on single cell sequencing (for losses and gains) of a standard breast cancer line in 2D culture that also exhibits a diversity of CNA clones similar to primary human breast tumors.^30^

Plotting ChReporter loss as a function of relative compressive strain *x* from the unperturbed height showed a good fit to a 2-state model (**Fig.1D**; R^2^ = 0.98). Similar mechanochemical models fit well to pressure-driven Piezo-1 opening^31^ and to curvature-driven Lamin-B dissociation from the nuclear envelope.^32^ Confinement-driven ChReporter loss could therefore have its origins in perturbation of a single, strained factor. The mitotic spindle is one possibility because of its known elongation, or it could also be a nuclear envelope factor given confinement-induced rupture of interphase nuclei.^11,33^

To begin to clarify key determinants of increased ChReporter loss after confinement, we assessed the sensitivity of A549 cells to known confinement-driven processes, starting with nuclear deformation. Confinement to 10µm did not alter nuclear area, but deeper confinement did increase nuclear area (∼5-fold for 2µm versus 19µm, **Fig.1E**). The nuclear periphery showed gaps in RFP-LMNB1 with GFP-H2B blebbing outward at rupture sites and a ∼30-fold increase in such ruptures for 2µm versus 2D (**Fig.1F**). We estimate ∼50% of nuclei ruptured in a 3µm confinement height, consistent with ∼50% nuclear rupture for migration through 3µm pores.^34^ Importantly, nuclear rupture *did not occur* for 10µm confinement relative to 2D (**Fig.1Fii**), which differs from increased ChReporter loss for 10µm. After 16h confinement, 6µm and 2D showed no difference in cell death whereas 2µm increased cell death, and no cells survived 64h of confinement (**Fig.1Fii-inset**). We increased nuclear rupture by LMNA knockdown^34^ but found no effect on ChReporter loss (**Fig.S1F**). Nuclear rupture increases DNA damage in these devices,^11^ but ChReporter loss clearly does not correlate with rupture (**Fig.1B-ii**), which argues against a role for DNA damage.

The lack of significant effect of 10µm confinement on interphase spreading and rupture is consistent with the typical interphase nuclear height of ∼10µm, which is much smaller than the height of mitotic chromatin (**Fig.1Aii**). We therefore hypothesized that confinement of 10µm or less perturbs mitosis, leading to ChReporter loss. To begin to assess this, we used a CDK4/6 inhibitor to block G1 exit and prevent progression through M phase during the standard confinement experiment. At the end of the experiment, no mitotic cells were detected, and ChReporter loss was greatly suppressed (**Fig.1G**). The latter functional results suggest that mitosis contributes to ChReporter loss, with further studies below showing the result relates to the cited need for 1-2 divisions that dilute and dim RFP-LMNB1 protein in cells with chromosome loss.^19^

Live cell imaging showed cells completing mitosis were *significantly decreased* for 10µm relative to 19µm (**Fig.1Hi,ii**). ChReporter losses were likewise high at 10µm but not 19µm versus 2D (**Fig.1Bii**). Strong confinement also suppressed mitotic events and prolonged pro-metaphase (∼10-fold for 4µm versus 10-20min for 19µm, **Fig.1Hi-iii,S1G**). The latter was expected.^14^ Intravital imaging of normal mouse skin epithelial stem cells in their basal niche also also show a pro-metaphase duration of 30-60min (**Fig.1Hiii-inset**), but estimates of chromatin heights from intensity and area across cell cycle stages suggest no significant compression of mitosis. Despite possible effects of cell type and species, mitotic compression *in vitro* generally lengthens the spindle versus 2D (**Fig.S1H**),^13,14^ and given that our MPS1i positive control for ChReporter loss is also a spindle-related perturbation, we focused on additional mitotic aberrations that are unstudied in confinement.

### Micronuclei that emerge only after mild confinement predict Chr loss

For 2µm confinement, imaging of mitotic cells after 16h of confinement showed pro/metaphase but no anaphase (versus 13% anaphase for 2D Ctrls), and chromosomes were splayed out (**Fig.2A**). Similar images of *intact* cells were shown previously^14^ and resemble standard metaphase spreads from ruptured cells where the Noc is often used to keep cells in pro/metaphase.^35^ For 6µm confinement, chromosomes are minimally splayed by the milder confinement on mitotic cells, but anaphase cells remained rare. Such images raise the possibility of mis-segregation of chromosomes into micronuclei, although there are no reports that mitotic confinement generates viable cells with micronuclei.

**Figure 2.**
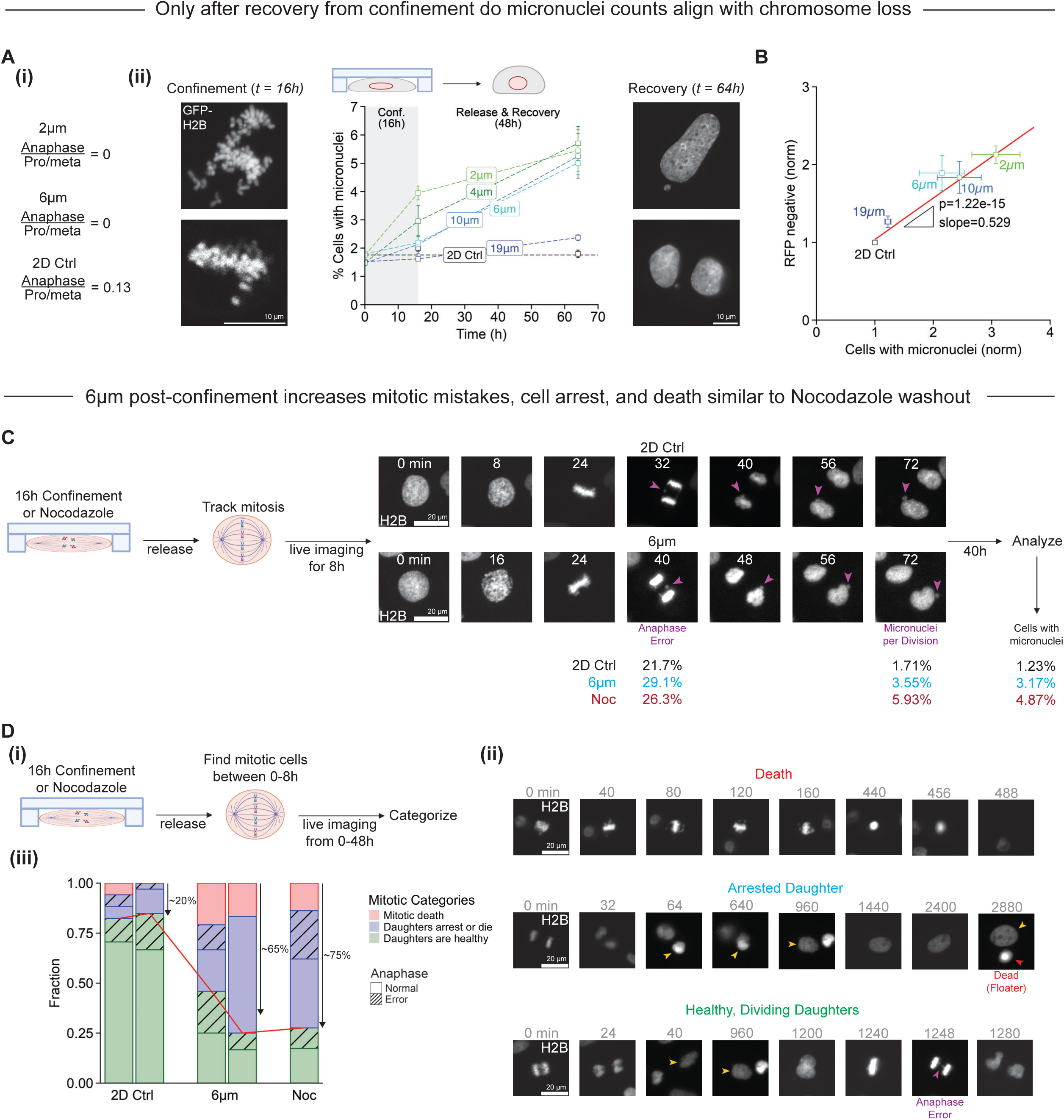
In recovery from confinement, mitotic chromosomes mis-segregate into micronuclei that track with ChReporter loss. Also see Figure S2. **(A)** Mitosis and micronuclei after confinement. **(i)** Quantification and **(ii)** images of GFP-H2B (i.e. H2B) pro/metaphase or anaphase cells at the end of 16h confinement to 2µm (n=18 cells), 6µm (n=22 cells), or 2D control (n=31 cells). Plot of micronuclei before confinement, at the end, and after recovery. (>450 cells per condition, n≥3 biological replicates; Mean, SEM). Scalebar = 10μm. **(B)** ChReporter loss versus micronuclei after recovery. Normalized to 2D control. (n≥3 biological replicates; Mean, SEM;_F-test). **(C)** Live image analyses of anaphase errors and micronuclei after release from 16h of 6µm confinement or 16h of Noc (0.1µM) (n>100 cells). Scalebar = 20μm. **(D)** Live image analyses of mitotic cells after release from 16h of 6µm confinement. **(i,ii)** Three fates: mitosis completes and daughters divide; mitosis is incomplete and cell dies; or mitosis completes but daughters arrest. **(iii)** Mitotic cells mostly die or arrest after confinement or washout of Noc (0.1µM), but death and arrest are rare for control (n≥29 cells). Scalebar = 20μm.

The 19µm confinement always maintained basal levels of ∼2% micronuclei similar to 2D Ctrls, but the 6µm and 10µm confinement started at baseline at 16h before increasing to ∼5% micronuclei only *after the 48h of recovery* (**Fig.2A**). The 2µm height showed 4% of cells had micronuclei already by the end of the 16h confinement, whereas 4µm results were between those for 2µm and 6µm. All confinement heights from 2-10µm nonetheless showed ∼5% micronuclei after 48h of recovery, and distributions of micronuclei diameters are independent of confinement height at mostly <3.5µm (**Fig.S2A**). Such a size suggests just 1-2 chromosomes within micronuclei.^25^ Given that chromosome loss typically results from micronucleus formation,^24^ we indeed find ChReporter loss associates with micronuclei *after* recovery from confinement (**Fig.2B**). The fitted slope and the unnormalized 2D control (∼0.1%) leads to an estimate that roughly 1 in every 40 micronuclei relates to loss of ‘tagged’ Chr-5. This seems reasonable given that A549 cells have ∼65 chromosomes.^18^ Our previous live-cell imaging study had shown some cells that generate micronuclei also lose the ChReporter.^19^

To directly show micronuclei arise from mis-segregation after mild confinement, we tracked live mitotic cells shortly after release (**Fig.2C**). Lagging chromosomes in anaphase could be seen to develop into micronuclei in one of the two daughter cells, which is the usual cause of immediate chromosome loss for one daughter and possible gain in the other.^24^ Such anaphase errors were 29.1% after confinement and only 21.7% for 2D controls. Nocodazole washout (after 16h) similarly increased anaphase errors to 26.3%, consistent with known effects of prolonged mitosis on chromosome cohesion.^21^ A fraction of these aberrant mitotic events led to micronuclei, which also increased from 1.7% in 2D control to 3.6% and 5.9% for 6µm confinement and nocodazole, respectively. Micronuclei trends remained the same after the full 48h recovery period, which aligns with quantitation for fixed samples (**Fig.2A**). These live-cell imaging differences begin to suggest that release from confinement is similar to widely used Noc washout in terms of causing aberrant mitosis and micronuclei that lead to Chromosome number changes.

Live-cell imaging was used to track the fate of any mitotic cell identified within 8h after confinement release – including the fates of daughter cells (**Fig.2Di-ii**). By the end of the 48h recovery period, we observed death or arrest in 65% of such cells compared to 20% for 2D controls (**Fig.2Diii**), and compared to a very low 3% death during confinement (6µm) (**Fig.1Fii-inset**). The high level of death or arrest after release from confinement also supports a ‘memory’ of mitotic confinement, as reported in past studies of Noc washout that showed for RPE1 cells with near-normal karyotype a similarly high degree of arrest or death of mitotic cells and daughters.^27,36^ Rescue of arrest by inhibition of both p38 and caspases was also documented, but we saw no such effects on ChReporter loss (**Fig.S2Bi**), and combining 6µm confinement with p38i also showed no significant difference (**Fig.S2Bii**). Compared to normal cells such as RPE1, one can expect cancer cells as used here have evolved different sensitivities and mechanisms in arrest and death under mitotic stress.

### Nocodazole washout phenocopies release from Confinement

To further compare confinement to Noc washout, we measured the kinetics of micronuclei generation and ChReporter loss. Immediately after washout, Noc-treated cells showed no increase in micronuclei counts versus 2D control (**Fig.3A**), which was expected as Noc prevents mitosis while prolonging pro/metaphase.^37^ At the early 16h and final 64h timepoints, the combination of Noc and 6µm confinement gave the same results as confinement or Noc alone, with micronuclei counts about 2-fold higher than 2D controls (**Fig.3A**). For Noc, the increase was expected.^26^ We chose 6µm height because ChReporter loss shows a small error bar and rupture and death are minimal (**Fig.1Biii**,**1F**). ChReporter loss at 48h (after recovery/washout) again associated with micronuclei counts (**Fig.S2D**), with each condition – separate or combined – showing ∼2-fold more loss relative to 2D control but not showing differences between each other (**Fig.3B**). Importantly, the lack of effect of the combination is consistent with spindle disruption being a key pathway regardless of the method. We confirmed this conclusion with an inhibitor of centrosome duplication that shortens spindle length during confinement:^38^ we observe increased ChReporter loss in 2D but no additional effect with confinement (**Fig.S2E**).

**Figure 3.**
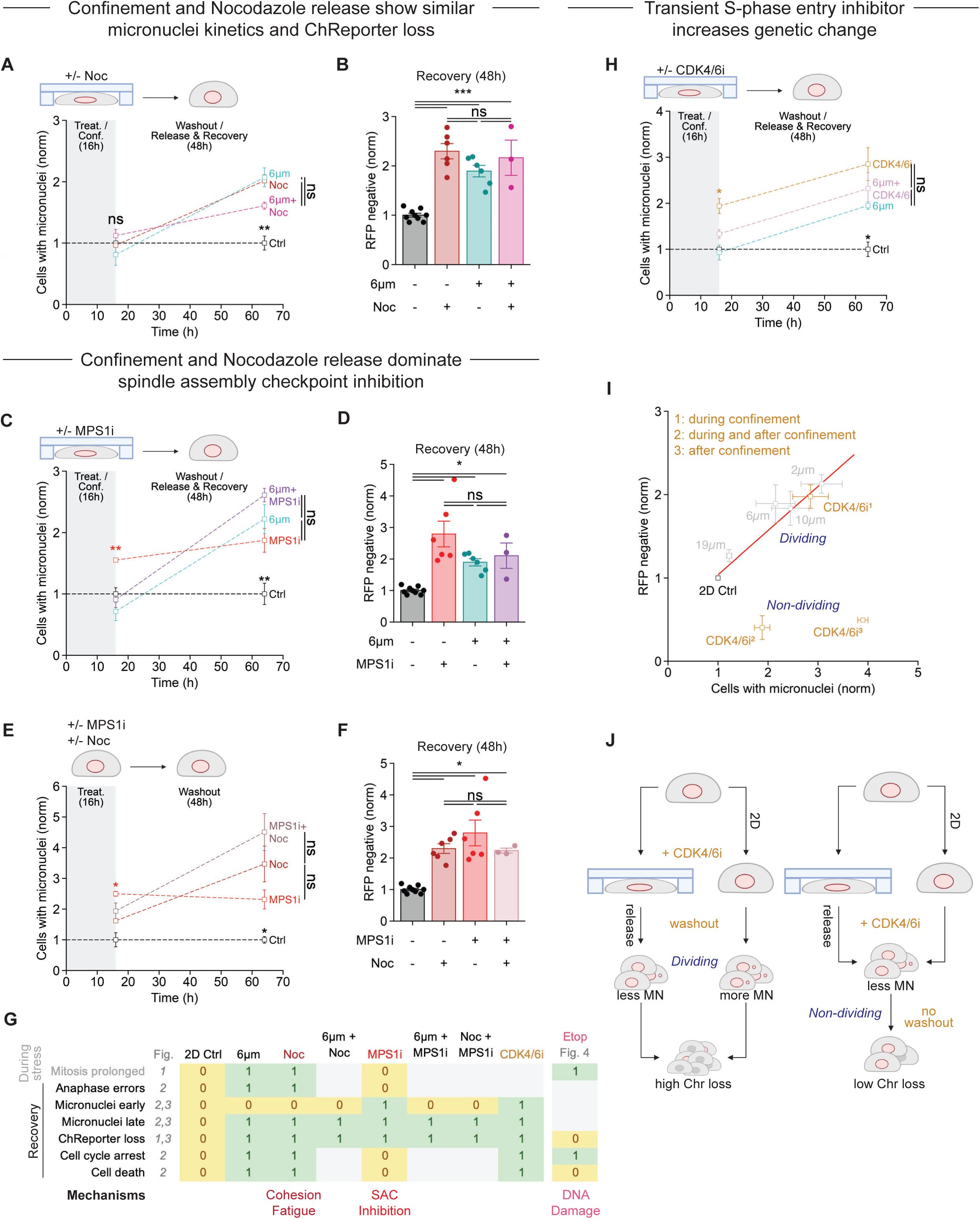
Inhibitors of spindle assembly checkpoint, microtubules, and CDK4/6 increase micronuclei and ChReporter loss in 2D but not moreso in confinement. Also see Figures S2 and S3, and Table S1. **(A)** Quantification of micronuclei for +/−Noc (0.1µM) at the end of +/−6µm confinement and after 2 days recovery normalized to control. (n=3 biological replicates; Mean, SEM; ordinary_one way_ANOVA_and_Tukey’s_multiple_comparisons_test; n.s. not significant, **p<0.01). **(B)** Flow cytometry of ChReporter loss for +/−nocodazole (0.1µM) and +/−6µm confinement normalized to 2D control. (n≥3 biological replicates; Mean, SEM; ordinary_one-way_ANOVA_and_Tukey’s_multiple_ comparisons_test; n.s. not significant, ***p<0.001). **(C)** Quantification of micronuclei for +/−MPS1i (50nM reversine) at the end of +/−6µm confinement and after 2 days recovery normalized to 2D control. (n=3 biological replicates; Mean, SEM; ordinary_one-way_ANOVA_and_Tukey’s_multiple_comparisons test; n.s. not significant, **p<0.01). **(D)** Flow cytometry analysis of ChReporter loss for +/−MPS1i (50nM reversine) and +/−6µm confinement normalized to 2D control. (n≥3 biological replicates from all drug experiments; Mean, SEM; ordinary_one-way_ANOVA and_Tukey’s_multiple_comparisons_test; n.s. not significant, *p<0.05). **(E)** Quantification of micronuclei for +/−MPS1i (50nM reversine) and +/−nocodazole (0.1µM) at the end of confinement and after 2 days recovery normalized to 2D control. (n=3 biological replicates; Mean, SEM; ordinary_one-way_ANOVA_and_Tukey’s_multiple_comparisons_test; n.s. not significant, *p<0.05). **(F)** Flow cytometry analysis of ChReporter loss for +/−MPS1i (50nM reversine, Rev) and +/−nocodazole (0.1µM) normalized to 2D control. (n≥3 biological replicates from all drug experiments; Mean, SEM; ordinary_one-way_ANOVA_and_Tukey’s_multiple_comparisons_test; n.s. not significant, *p<0.05). **(G)** Confinement and transient drug summary. Noc washout phenocopies confinement. Proliferation seems unaffected by Rev, and death does not increase from 2D Ctrl for Rev (∼0.8-0.9%) but does increase to ∼2% for CDK4/6i. **(H)** Quantification of micronuclei for +/−CDK4/6i (10µM Palbociclib) at the end of +/−6µm confinement and following 2 days recovery normalized to 2D control. (n=3 biological replicates; Mean, SEM; ordinary_one-way_ANOVA_and_Tukey’s_multiple_comparisons test; n.s. not significant, *p<0.05). **(I)** ChReporter loss versus micronuclei seen after recovery. Normalized to 2D control. (n≥3; Mean, SEM). **(J)** Summary of CDK4/6i treatments and confinement. Treatment during confinement: high micronuclei counts and ChReporter loss. Treatment after confinement: high micronuclei counts but low ChReporter loss. Division during recovery from treatment leads to similar micronuclei and ChReporter trends.

### Mild confinement suppresses micronuclei caused by MPS1i

MPS1i treatment for just 16h induced a significant increase in micronuclei above basal levels (**Fig.3C**), consistent with known effects.^28^ Surprisingly, MPS1i’s acute induction of micronuclei was suppressed to control levels when combined with 6µm confinement. Micronuclei levels at 64h were non-additive when combined with 6µm confinement. Regardless, the non-additivity with MPS1i is consistent with a confinement-induced perturbation of the spindle that also impacts the spindle assembly checkpoint. We tested this hypothesis by *chemically* perturbing the *confined* spindle. By 48h after confinement, all conditions (MPS1i, confinement, and the combined) resulted in ∼2-fold increases in micronuclei and ChReporter loss versus 2D controls (**Fig.3D**,**S2C**). Importantly, the 48h result is the same *endpoint* as Noc despite the difference at 16h (**Fig.3A**). These results are consistent with MPS1i and confinement ultimately targeting the same or a related pathway in chromosome loss that correlates with micronuclei induction **(Fig.2B)**. We also rule out pathway saturation as higher drug doses were tested earlier.^19^ In contrast, myosin-II inhibition during mild compression caused more ChReporter loss of the same A549 cells and other cell types.^19^ Hence synergistic or complementary pathways certainly do exist.

Whereas micronuclei are increased by MPS1i after the 16h treatment and remain the same 48h later, the combination of Noc + MPS1i at release does not differ from 2D control and has increased 48h later (**Fig.3E**). The trends are the same for 6µm confinement + MPS1i (**Fig.3C**). Moreover, after the 48h recovery, all treatments again showed similar 2-3 fold higher levels of micronuclei and ChReporter loss relative to controls (**Fig.3E,F**; **Table_S1**). Overall, these results further indicate that confinement is phenocopied by Noc and not by a spindle assembly checkpoint disruption (**Fig.3G**).

### Transiently blocking cell cycle entry with CDK4/6i increases Chromosome loss

Given mitotic perturbations associate with ChReporter loss after confinement, we transiently inhibited cell cycle entry with CDK4/6 inhibition (CDK4/6i) *only during* the 16h confinement. Micronuclei are generated by CDK4/6i,^29^ and we find the trends align with MPS1i results for micronuclei kinetics as well as ChReporter loss whether combined or not with 6µm confinement (compare **Fig.3H,S3A** and **3E,F**). Mitotic counts after the 48h CDK4/6i recovery period (without confinement) were similar to those of 2D control, and we also noted increased nuclear area following treatment (**Fig.S3A**). Combining CDK4/6i with MPS1i also increased micronuclei levels at 16h and persisted but were not significantly different from each drug alone (**Fig.S3B**), which suggests related pathways.

Following up on our initial studies with sustained CDK4/6i treatment during confinement (**Fig.1G**), we also treated cells with CDK4/6i only after 6µm confinement. Cell cycle progression during the 48h recovery was thus blocked in both approaches, with no mitosis observed at the end of recovery (**Fig.S3C-H**). ChReporter loss decreased relative to 2D control with drug alone and when combined with confinement even though micronuclei counts were once again elevated. This apparent paradox is resolved by recalling that loss of the ChReporter absolutely requires cell divisions to dilute and dim the RFP-LMNB1 protein (**per Fig.1A,G**) – as directly visualized in our initial studies.^19^ Thus we conclude that high micronuclei counts are *necessary but not sufficient* for ChReporter loss as further division is also needed for loss (**Fig.3I,J**).

### Transcriptional impact of confinement shows Chr loss and dysregulated mitosis

Given our focus on the very rare cells that are ChReporter-negative (∼1-per-500 cells in **Fig.1B-i**), we hypothesized that chromosome loss associates with persistent changes in the transcriptome – particularly decreased expression from Chr-5. Therefore, we sorted live cells for ChReporter loss after the 48h recovery from 6µm confinement or 2D controls, and then we used single-cell RNA-seq (scRNAseq) to quantify transcript changes (**Fig.4A, S4A**). Relative to RFP-positive control cells as a reference (see inferCNV in Methods), the sorted RFP-negative cells showed most Chr-5 transcripts were down-regulated for 6µm confinement or 2D controls (**Fig.4B**). Some cells also showed evidence of rare gains and losses in other chromosomes (e.g. loss of Chr-3, gain of Chr-12, gains and losses of Chr-20), which either existed before confinement or were induced. The spontaneous RFP-negative cells from 2D cultures largely segregated from the confined cells along both UMAP1 and UMAP2 axes (**Fig.4C-left**). The oncotherapy-relevant mitosis gene *TOP2A* was upregulated >3-fold in most of the confined cluster and also in a minor cluster of control cells nearby (**Fig.4C-right**, **S4A**). Because *TOP2A* levels increase with UMAP1, we anticipated that the most enriched gene ontology (GO) terms relate to mitosis such as ‘Chromosome segregation’ (**Fig.4D**,**_Table_S3**).

**Figure 4.**
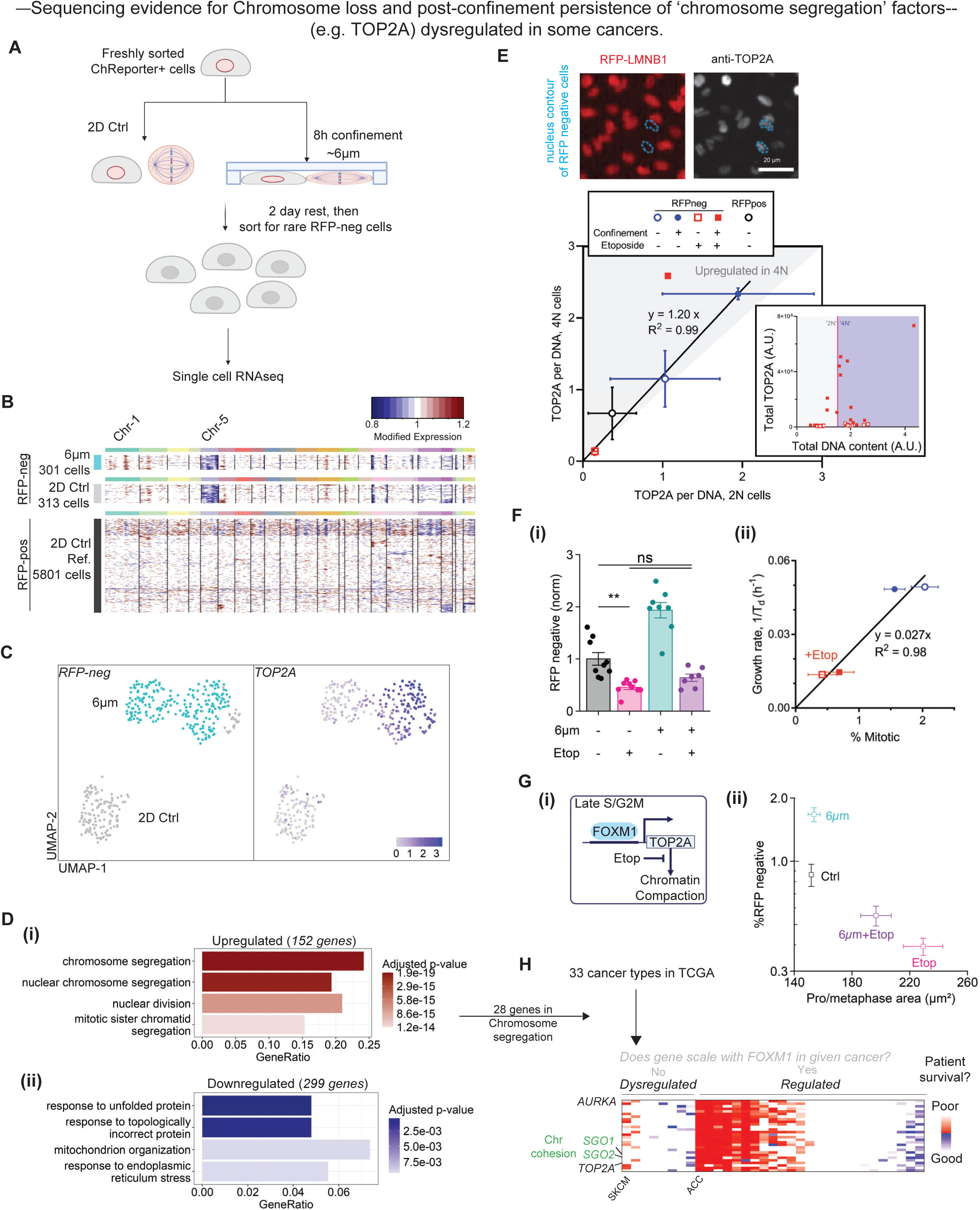
Post-confinement persistence of ‘chromosome segregation’ genes and their dysregulation in some cancers. Also see Figure S4 and Tables S3,4. **(A)** Experimental design for single-cell RNA-sequencing of 6µm confined cells. Freshly sorted ChReporter-positive A549 cells are confined for 8h. After 2 days of rest, both confined and 2D control cells are sorted for ChReporter-negative cells and then sequenced. **(B)** Heatmaps from inferCNV show loss of Chr-5 ChReporter and decreased transcripts. **(C)** UMAP of ChReporter-negative cells. Each point is a cell. Cluster colors are for confined or 2D cells. **(D)** Gene ontology GO enrichment analysis of differentially expressed genes in confined versus unconfined groups. Genes with p< 0.05 and a log2FoldChange > 1 were used in analysis. **(i)** Upregulated ontologies relate to division and chromosome segregation. **(ii)** Downregulated ontologies relate to endoplasmic reticulum stress and unfolded protein response. **(E)** Image: ChReporter cells after 6µm confinement (8h with 48h recovery) were stained with anti-TOP2A. Blue outline indicates nucleus contour of RFP-negative cells as obtained from DNA stain (Hoechst 33342). Plot: Quantification of fluorescence intensity of anti-TOP2A per fluorescence intensity of DNA from 4N cells (late-S/G2/M) versus 2N cells (G1/early S). Etop (1µM) was used during the confinement period. (inset plot) Example identification of 4N and 2N cells based on total fluorescence intensity for DNA. (>20cells for n≥3 biological replicates; Mean, SEM; ordinary_one-way ANOVA and Tukey’s multiple comparisons_test; ****p<0.0001). Scalebar = 20μm. **(F)** Flow cytometry of ChReporter loss for +/−Etop (1µM) and +/−6µm confinement normalized to 2D control. **(inset)** Growth rate versus mitotic cell counts. (n≥3 biological replicates; Mean, SEM; ordinary_one-way_ANOVA_and_Tukey’s_multiple_comparisons test; n.s. not significant, **p<0.01). **(G)** ChReporter loss versus pro/metaphase area for +/−Etop (1µM) and +/−6µm confinement. (n≥3 biological replicates; Mean, SEM; ordinary_one-way_ANOVA_and_Tukey’s_multiple_comparisons_test; n.s. not significant, *p<0.05). **(H)** Schematic for patient prognosis heatmaps based on high and low expression of mitotic genes.

To assess whether gene expression changes observed are part of a general confinement signature irrespective of whether cells have lost one copy of chromosome 5 or not, we applied bulk RNA-seq to all cells after confinement versus 2D controls. For further generality, we used strong confinement of 2µm and breast ductal carcinoma in situ (DCIS) cells (**Fig.1Bii**) because the growth of such cells is already known to be affected by confinement within the mammary duct.^11^ Cells were confined for 2h, with some mitotic cells evident. Surprisingly, the deep sequencing shows over half of the top-25 enriched GO terms related to chromosome segregation and mitosis versus 2D controls (**Table_S4**), consistent with scRNAseq of rare ChReporter-negative cells (**Fig.4D**).

Bulk RNA-seq gene sets included *FOXM1*, which is a master regulator transcription factor for many mitotic genes including *TOP2A* and *LMNB1* (**Fig.S4Bi-iv**).^39^ Interestingly, downregulated GO terms in the scRNA-seq and bulk RNAseq datasets did not overlap (**Fig.4D, Table_S4**). This might be because 2µm causes much more strain on chromatin (**Fig.1E,F,2A-ii**), leading to more severe damage and cell death (**Fig.1Fii-inset**). Accordingly, downregulated GO terms under 2µm confinement are metabolism-related, whereas those under 6µm confinement involve protein unfolding and organization pathways.

Protein levels of TOP2A were upregulated following a 6µm confinement with a 48h rest, consistent with mRNA results, and protein was elevated not only in late cell cycle as expected but also unexpectedly in early cell cycle (**Fig.4E,S4A,Biv**). Cell division is blocked by etoposide (Etop) which inhibits TOP2A, causing DNA damage. Given that chromosome loss requires cell division (**Fig.1A**)^19^ as confirmed by sustained CDK4/6i (**Fig.1G,S3C,D**), low dose Etop indeed suppressed ChReporter loss and proliferation, although the %RFP-negative cells trended higher for confined than in 2D Etop controls (**Fig.4F**). The high TOP2A that results from confinement (**Fig.4C,E**) could be less inhibited by Etop: mitotic compaction is indeed greater than seen for Etop in 2D (**Fig.4Gi,ii**), consistent with TOP2A-driven compaction of mitotic chromatin.^40,41^ Regardless, transient treatment with DNA-damaging Etop does not phenocopy the effect of confinement (**Fig.3G**).

## DISCUSSION

Release from both mild confinement and Noc increases mis-segregation into micronuclei with similar kinetics (**Fig.2C,3G**), with mechanisms for Noc washout including fatigue of the cohesion between sister chromatids and an end-result being aneuploidy.^21,42,43^ Release from both mild confinement and Noc also cause daughter cell arrest and death (**Fig.2D,3G**). Such arrest and death are one facet of the ‘*memories*’ of prolonged mitosis.^27,36^ In contrast, MPS1i and CDK4/6i both generate micronuclei with distinctly rapid kinetics – unless the cells are confined. The results thus suggest a multi-faceted memory of prolonged mitosis in confinement that extends to mis-segregation into micronuclei and heritable chromosome loss.

Clinical trials of CDK4/6i’s against breast cancer show relapse is common.^44,45^ Micronuclei are known to be generated by CDK4/6i.^29^ Our results suggest micronuclei will increase genetic heterogeneity that can fuel resistance to therapy, but how CDK4/6i functions during confinement is an important question, given the confinement of breast cancer cells *in vivo*.^11^ At least for 2D cultures, we speculate that cells which are already in S/G2/M when suddenly exposed to this clinical inhibitor will mis-segregate chromosomes into micronuclei and then the cells will stop in G0/G1. Further studies of kinetics should address such mechanisms.

Past observations of daughter cell arrest after prolonged mitosis did not study micronuclei or ‘chromosome segregation’ pathways,^27^ but we observe that this is a top GO term upregulated by confinement in both scRNAseq (30 genes) and bulkRNAseq (409 genes) datasets (**Fig.4Di**, **Table_S4**). We find 27 genes overlap (**Fig.S4Bi**), and we added the master transcription factor of cell cycle *FOXM1* (to make 28 genes) as it is likely too low for scRNAseq detection (**Fig.S4C**). Chromosome cohesion genes *SGO1* and *SGO2* are among the 28 genes, which indicates the confined cells are upregulating cohesion that is impaired in prolonged mitosis.^21^ Since only a fraction of mitotic genes are persistently upregulated by confinement (24% for ‘chromosome segregation’, **Fig.4Di**), the mitotic program seems dysregulated. Therefore we analyzed 33 cancers in The Cancer Genome Altas (TCGA) for correlations between the key mitotic genes, and we also assessed patient survival (**Fig.4H**). We first quantified correlations between *FOXM1* and its known target *TOP2A* (**Fig.4G-i**) for all cancer types (**Fig.S4Ci-ii**) and find some cancers lack the usual correlation and are thus ‘dysregulated’ (**Fig.S4C-iii**). For nearly all such cancers, patient survival and *TOP2A* levels do not correlate, whereas high *TOP2A* in the ‘regulated’ cancers often associates with poor survival (**Fig.S4C-iv**). The trend holds across the 28 genes, with predictive value of only 8.5±1.9% for dysregulated cancers versus 38.3±1.8% for regulated cancers (*p<*0.05). One possible scenario is that *TOP2A* stays high (**Fig.4C**) in arrested daughter cells (**Fig.2D**) with suppression of other mitotic genes. Stress-driven dysregulation of mitotic genes might thus decouple standard expression signatures from clinical outcomes and might serve as a marker for cell confinement.

### Limitations of the study

Our ChReporter system tracks protein loss, but loss of tumor suppressor proteins subsequent to chromosome loss is prevalent in the aneuploidy of cancers. Our ChReporter can in principle detect gains as bright cells, which is more relevant to possible fates of a micronucleus in one of the two daughter cells. Identifying expression signatures (e.g. **Fig.4**) that associate with confinement and that might even relate to heritable mutations in a first stage of mechano-evolution requires additional study. However, expression signatures of confinement might explain some aspects of human tumor expression data, such as a correlation between chromosome losses or gains with poor survival rates in most cancers including lung adenocarcinoma (LUAD).^46^

Confinement times *in vivo* are unknown, and while cells can survive 16h of our *in vitro* confinement, longer times *in vitro* increase cell death. Formation of micronuclei as a consequence of confinement had not been reported previously and occurs after release as does release from CDK4/6 inhibition (**Fig.3H**). In order to more rigorously demonstrate heritable loss at a single cell level, we must track many cells for multiple days so as to observe a chromosome mis-segregate into a micronucleus followed by loss of cell fluorescence and generation of a colony. This type of experiment requires about a week of continuous imaging, along with some luck, to capture the rare events, but it could add important insight.

## RESOURCE AVAILABILITY

### Lead contact

Requests for further information and resources should be directed to the lead contact, Dennis E. Discher

### Materials availability

This study did not generate new unique reagents. There are restrictions to the availability of cell lines and confinement devices used here because of the lack of an external centralized repository for their distribution. We are glad to share with reasonable compensation by requestor for its processing and shipping.

### Data and code availability

- All data reported in this paper will be shared by the lead contact upon request.
- This paper does not report original code.
- Single-cell RNA-seq data have been deposited at GEO and are publicly available as of the date of publication. Accession numbers are listed in the key resource stable.
- Any additional information required to reanalyze the data reported in this paper is available from the lead contact upon request.

## ACKNOWLEDGMENTS

We are grateful to The Penn Cytomics and Cell Sorting Resource Laboratory and The Penn Genomic and Sequencing Core. We are also grateful for technical assistance of former Discher Lab members at Penn Brandon Hayes and Peter Zhu as well as Xinran Ren (for analysis of the micronuclei diameters in the 2μm condition), Dr. Manu Tewari, and Joanna S. Georgiou for valuable comments on the manuscript. We also thank members of the Nader Lab for their input: Diana I. Cruz, Akash Samuel Carasala, Logan Northcutt. This work was supported by funding from the following sources: NIH U01 CA254886 (DED), P01 CA265794 (DED), National Science Foundation CMMI1548571,154857.

## AUTHOR CONTRIBUTIONS

Conceptualization, D.E.D.; methodology, S.H.P., M.W., P.R., G.N., D.E.D.; Investigation, S.H.P., M.W., K.R.S., J.K., J.S.D., P.R.; writing—original draft, S.H.P., D.E.D.; writing—review & editing, S.H.P., G.N., D.E.D.; funding acquisition, D.E.D.; resources, G.N. and D.E.D.; supervision, G.N. and D.E.D.

## DECLARATION OF GENERATIVE AI AND AI-ASSISTED TECHNOLOGIES

AI was not used

## SUPPLEMENTAL INFORMATION

### Supplemental Figure & Table Legends

**Figure S1.**
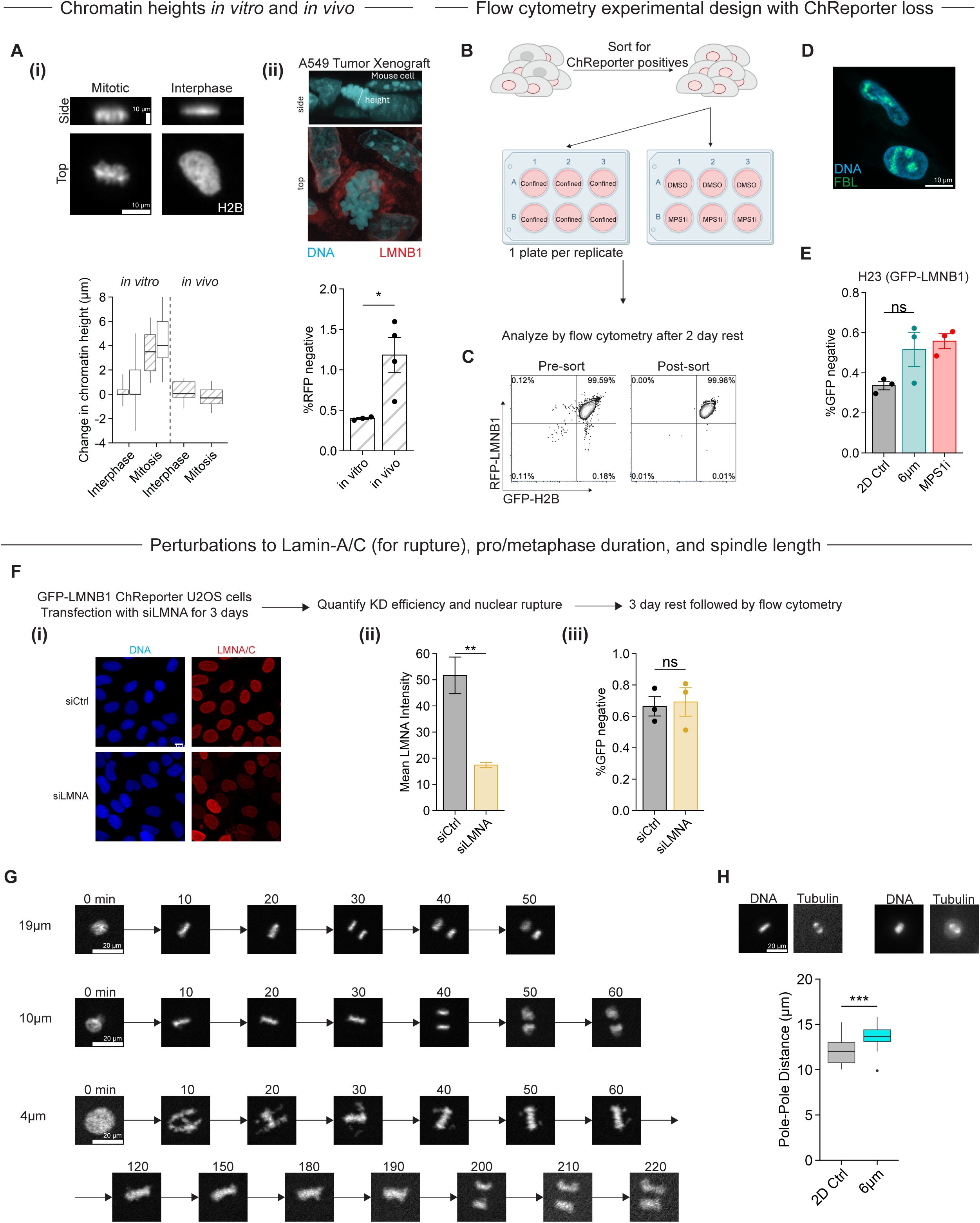
Tumor-relevant confinement induces genetic change in another lung adenocarcinoma line, but knockdown of LMNA does not. Related to Figure 1. **(A)** Chromatin images and height differences relative to interphase *in vitro* and *in vivo.* **(i,ii)** Chromatin heights relative to interphase in standard 2D cultures or for tumors *in vivo.* Measurements are reported as a difference relative to 2D-interphase because of different imaging systems and resolutions that affect absolute measurements but not the differentials. The representative image shows RFP-LMNB1 and DNA stain for A549 tumor xenograft (scalebar for “height” = 10mm). Cross-hatched results determined from [Hayes 2023] (n≥30 cells per condition; the box represents the 25–75th percentiles, and the median is indicated; the whiskers show the 5–95% range) ChReporter changes for A549 cells are also from [Hayes 2023]. **(B)** Schematic detailing the protocol for cell confinement. Cells are first sorted for a pure (99.98%) population of ChReporter positive cells. For confined conditions, 1 plate is pooled together to be considered a technical replicate. For unconfined conditions, a single well is a replicate. Replicates are then analyzed by flow cytometry for ChReporter negative cells. **(C)** Representative flow cytometry plots of pre- & post-sort of A549 Chr-5 ChReporter (RFP-LMNB1) cells. **(D)** Image of A549 Chr-19 ChReporter (GFP-FBL) cells. **(E)** Quantification of H23 Chr-5 ChReporter (GFP-LMNB1) loss following 6µm confinement. (n=3; Mean, SEM; ordinary one-way ANOVA and Tukey’s multiple comparisons test; n.s. not significant). **(F)** Analysis of LMNA knockdown in U2OS Chr-5 reporter cells (GFP-LMNB1). **(i)** Images of U2OS cells following siLMNA or siCtrl after 3 days. **(ii)** There is ∼60% reduction in LMNA intensity. (n≥3; Mean, SEM; ordinary one-way ANOVA and Tukey’s multiple comparisons test; **p<0.01). **(iii)** Quantification of ChReporter loss after knockdown of LMNA. (n=3 replicates; Mean, SEM; ordinary one-way ANOVA and Tukey’s multiple comparisons test; n.s. not significant). **(G)** Time series of an A549 cell undergoing mitosis while confined to 19µm, 10µm, and 4µm, respectively. **(H)** Quantification of pole-pole distance of mitotic cells following 6µm confinement. (n≥10 cells; Mean, SEM; ordinary one-way ANOVA and Tukey’s multiple comparisons test; ***p<0.001).

**Figure S2.**
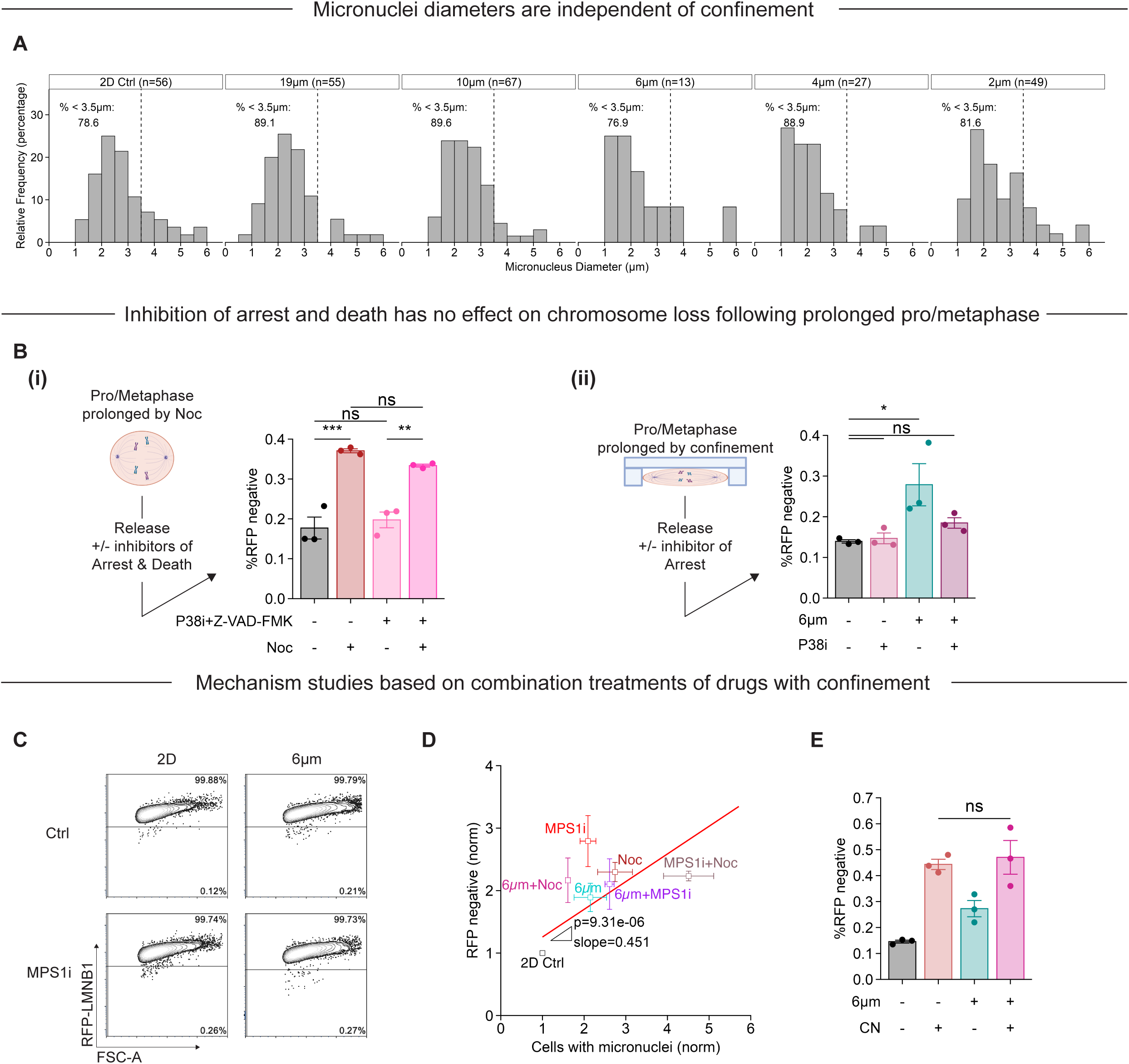
Inhibition of arrest and death has no effect on chromosome loss following prolonged pro/metaphase. Related to Figures 2 and 3. **(A)** Histograms of micronuclei diameters for 2D control cells and cells confined to 19µm (n=55 cells), 10µm (n=67 cells), 6µm (n=13 cells), 4µm (n=27 cells), and 2µm (n=49 cells). **(B)** Rescue of arrest and death following prolonged pro/metaphase. **(i)** Cells are treated with 0.3µM nocodazole. After washout, cells are treated with 10µM p38i and 20μM pan-caspase inhibitor (Z-VAD-FMK) for 8h. An additional 40h of rest if given before analysis. Quantification of ChReporter loss. (n=3; Mean, SEM; ordinary one-way ANOVA and Tukey’s multiple comparisons test; n.s. significant, **p<0.01, ***p<0.001). **(ii)** Cells are confined to 6µm for 16h. After release, cells are treated with 10µM p38i. Quantification of ChReporter loss. (n=3; Mean, SEM; ordinary one-way ANOVA and Tukey’s multiple comparisons test; n.s. not significant, *p<0.05). **(C)** Representative flow cytometry plots for +/− MPS1i and +/− 6µm confinement. **(D)** ChReporter loss versus micronuclei seen after recovery. Normalized to 2D control. (n≥3; Mean, SEM; F-test). **(E)** ChReporter loss for +/−centrinone (0.125µM) and +/−6µm confinement normalized to 2D control. (n=3; Mean, SEM; ordinary one way ANOVA and Tukey’s multiple comparisons test; n.s. not significant).

**Figure S3.**
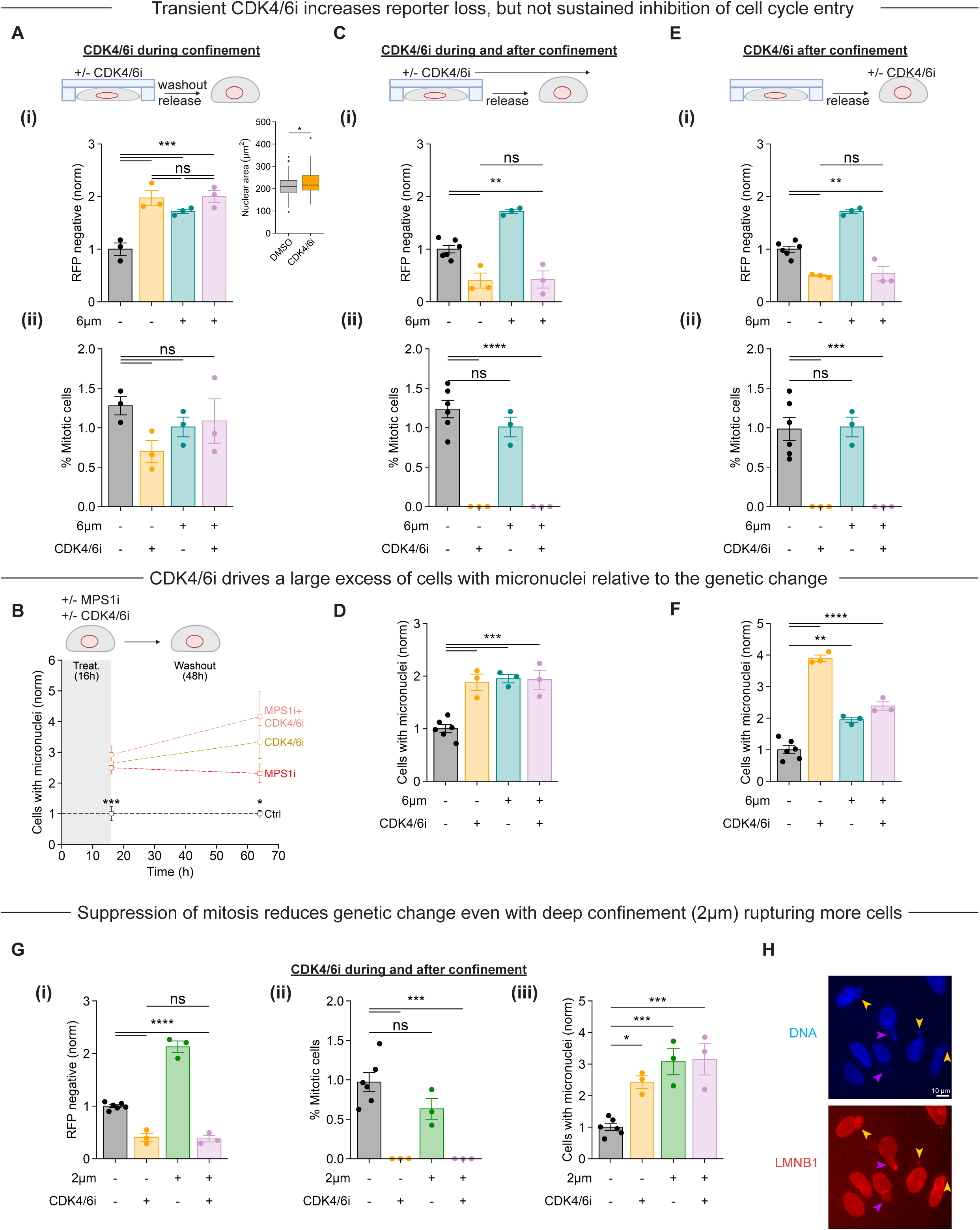
Inhibition of mitosis by CDK4/6i *after* confinement suppresses reporter loss, even though CDK4/6i induces micronuclei. Related to Figure 3. **(A)** CDK4/6i is added during confinement (16h) and washed out, allowing recovery. **(i)** Quantification of ChReporter loss for +/− CDK4/6i (10µM) and +/− 6µm confinement normalized to 2D control. (n=3; Mean, SEM; ordinary one-way ANOVA and Tukey’s multiple comparisons test; n.s. not significant, ***p<0.001). **(inset)** Quantification of nuclear area following 16h CDK4/6i (10µM). (n=100 cells; Mean, SEM; ordinary one-way ANOVA and Tukey’s multiple comparisons test; *p<0.05). **(ii)** Quantification of mitotic cells. (n=3; Mean, SEM; ordinary one-way ANOVA and Tukey’s multiple comparisons test; n.s. not significant). **(B)** Quantification of micronuclei for +/− MPS1i (50nM reversine) and +/− CDK4/6i (10µM) at the end of confinement and after 2 days recovery normalized to 2D control. (n=3; Mean, SEM; ordinary one-way ANOVA and Tukey’s multiple comparisons test; *p<0.05, ***p<0.001). **(C)** CDK4/6i treatment is sustained both during (16h) and after (48h) confinement. **(i)** Quantification of ChReporter loss for +/− CDK4/6i (10µM) and +/− 6µm confinement normalized to 2D control. (n≥3 from all drug experiments; Mean, SEM; ordinary one-way ANOVA and Tukey’s multiple comparisons test; n.s. not significant, **p<0.01). **(ii)** Quantification of mitotic cells. (n≥3 from all drug experiments; Mean, SEM; ordinary one-way ANOVA and Tukey’s multiple comparisons test; n.s. not significant, ****p<0.0001). **(D)** CDK4/6i is added after confinement (48h). **(i)** Quantification of ChReporter loss for +/− CDK4/6i (10µM) and +/− 6µm confinement normalized to 2D control. (n≥3 from all drug experiments; Mean, SEM; ordinary one-way ANOVA and Tukey’s multiple comparisons test; n.s. not significant, **p<0.01). **(ii)** Quantification of mitotic cells. (n≥3 from all drug experiments; Mean, SEM; ordinary one-way ANOVA and Tukey’s multiple comparisons test; n.s. not significant, ***p<0.001). **(E)** Quantification of micronuclei for +/− CDK4/6i (10µM) and +/− 6µm confinement. CDK4/6i treatment is sustained both during (16h) and after (48h) confinement. (n≥3 from all drug experiments; Mean, SEM; ordinary one-way ANOVA and Tukey’s multiple comparisons test; ***p<0.001). **(F)** Quantification of micronuclei for +/− CDK4/6i (10µM) and +/− 6µm confinement. CDK4/6i is added after confinement. (n≥3 from all drug experiments; Mean, SEM; ordinary one-way ANOVA and Tukey’s multiple comparisons test; **p<0.01, ****p<0.0001). **(G)** CDK4/6i is maintained both during (16h) and after (48h) 2µm confinement. **(i)** Quantification of ChReporter loss for +/− CDK4/6i (10µM) and +/− 2µm confinement normalized to 2D control. (n≥3 from confinement experiments; Mean, SEM; ordinary one-way ANOVA and Tukey’s multiple comparisons test; n.s. not significant, ****p<0.0001). **(ii)** Quantification of mitotic cells. (n≥3 from confinement experiments; Mean, SEM; ordinary one-way ANOVA and Tukey’s multiple comparisons test; n.s. not significant, ***p<0.001). **(iii)** Quantification of micronuclei for +/− CDK4/6i (10µM) and +/− 2µm confinement. (n≥3 from confinement experiments; Mean, SEM; ordinary one-way ANOVA and Tukey’s multiple comparisons test; *p<0.05, ***p<0.001). **(H)** Images of A549 cells following 16h CDK4/6i (10µM) treatment. Orange arrows point to micronuclei and magenta arrows point to blebs.

**Figure S4.**
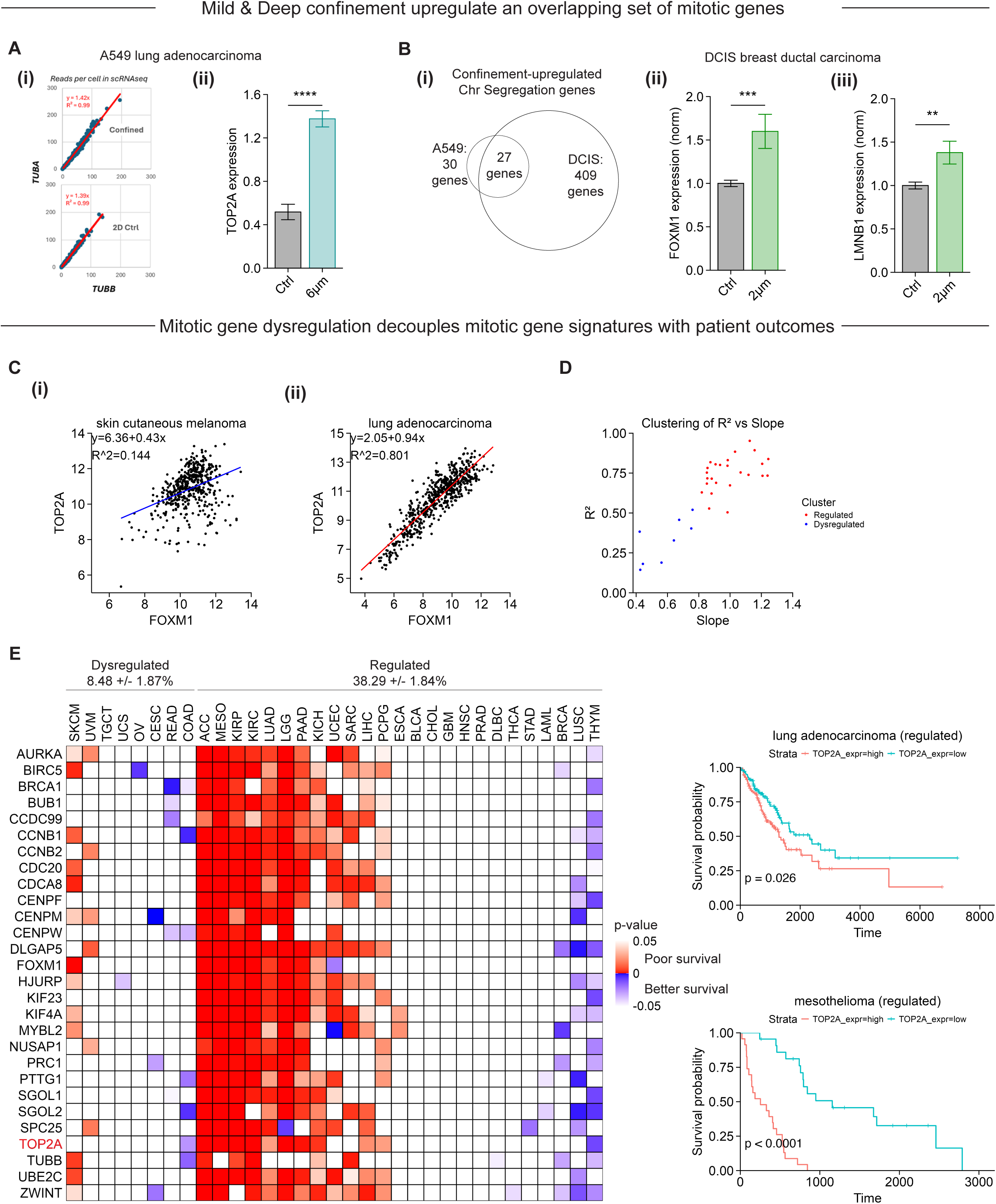
Confinement induced chromosome segregation gene signatures indicate dysregulation in some cancers lack correlations with survival. Related to Figure 4. **(A) (i)** Internal control validation for scRNAseq of A549 study: reads per cell for housekeeping genes *TUBA* (*A1B* + *A1C*) and *TUBB* (*B* + *B4B*). Linearity is expected for microtubule assembly of TUBA and TUBB heterodimers. **(ii)** Normalized TOP2A expression of 6µm confined cells from scRNA-seq data. (n≥100 cells; Mean, SEM; ordinary one-way ANOVA and Tukey’s multiple comparisons test; ****p<0.0001). **(B)** Bulk RNA-seq of 2µm confined DCIS cells. **(i)** Overlap of upregulated genes from ‘chromosome segregation’ gene set between scRNA-seq and bulk RNA-seq. The overlap set plus FOXM1 is listed in panel-(C). Normalized expression from bulk RNA-seq of **(ii)** *FOXM1*, and **(iii)** *LMNB1*. (Mean, SEM; n.s. not significant, **p<0.01, ***p<0.001). **(C)** *TOP2A* versus *FOXM1* expression from Pan-Cancer TCGA dataset for **(i)** melanoma (SKCM) and **(ii)** lung adenocarcinoma (LUAD). TOP2A and other mitotic genes are highly upregulated during G2/M by the transcription factor FOXM1 [Vashisth 2021], but a pro/metaphase extended by even 2-fold is expected to drive a corresponding 2-fold increase in TOP2A and other G2/M genes at both RNA and protein levels [Fischer 2022], consistent with results observed here (Fig.4E). Temporal integration of protein expression across mother-daughter generations, influenced by translation during different cell cycle phases, has been documented for mitogens [Min 2020], and confinement can have a similar effect on gene and protein regulation (Fig.4). **(D)** Plotting fits for all cancer types (R^2^ versus slope), cancer types were clustered based on poor or good regulation. **(E)** Patient prognosis heatmap based on expression of mitotic genes. Log-rank p-values based on Kaplan-Meier curves generated for each gene and each cancer type (examples for LUAD and MESO shown), based on the top (red) and bottom (blue) quartiles of expression. In the heatmap, red indicates high expression of gene leads to poor prognosis. Blue indicates high expression of gene leads to improved prognosis. White indicates no significance.

**Table S1.**
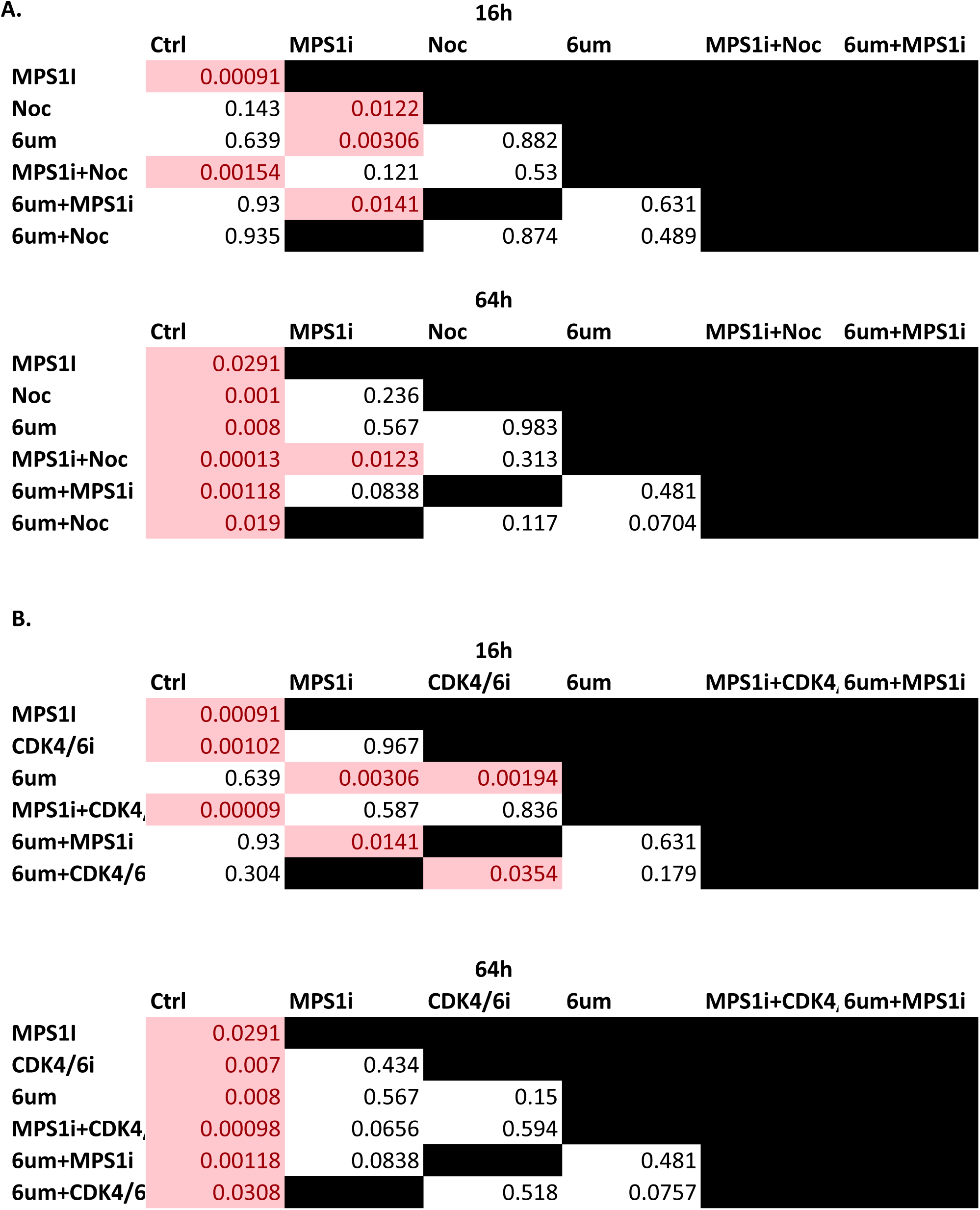
Matrices of p-values for MPS1i, nocodazole, CDK4/6i, and confinement results. Related to Figures 2 and 3. **(A)** Results for combinations of MPS1i, nocodazole, and confinement. **(B)** Results for combinations of MPS1i, CDK4/6i, and confinement. For comparisons with multiple p-values (coming from multiple experiments), the geometric mean is used. Boxes shaded red have p<0.05. (Ordinary one-way ANOVA and Tukey’s multiple comparisons test).

**Table S3.** Expanded list of gene ontology enrichment analysis. Related to Figure 4. The top 10 significantly upregulated and downregulated gene ontologies are shown.

| GENE ONTOLOGY NAME | GENE RATIO | ADJ p-val |
| --- | --- | --- |
| chromosome segregation | 0.241935484 | 1.89E-19 |
| nuclear chromosome segregation | 0.193548387 | 7.35E-16 |
| nuclear division | 0.209677419 | 7.39E-15 |
| mitotic sister chromatid segregation | 0.153225806 | 1.15E-14 |
| sister chromatid segregation | 0.161290323 | 2.02E-14 |
| mitotic nuclear division | 0.169354839 | 4.02E-14 |
| regulation of mitotic sister chromatid separation | 0.096774194 | 1.36E-12 |
| mitotic sister chromatid separation | 0.096774194 | 2.18E-12 |
| mitotic cell cycle phase transition | 0.185483871 | 6.57E-12 |
| regulation of mitotic nuclear division | 0.112903226 | 8.46E-12 |
| response to unfolded protein | 0.04797048 | 0.00045942 |
| response to topologically incorrect protein | 0.04797048 | 0.00084825 |
| mitochondrion organization | 0.073800738 | 0.00919489 |
| response to endoplasmic reticulum stress | 0.055350554 | 0.00926688 |
| endoplasmic reticulum unfolded protein response | 0.029520295 | 0.01825307 |
| regulation of mitochondrion organization | 0.033210332 | 0.01825307 |
| Golgi vesicle transport | 0.055350554 | 0.01825307 |
| purine ribonucleoside triphosphate metabolic process | 0.04797048 | 0.02576394 |
| mitochondrial transport | 0.040590406 | 0.02576394 |
| cellular response to unfolded protein | 0.029520295 | 0.02576394 |

**Table S4.**
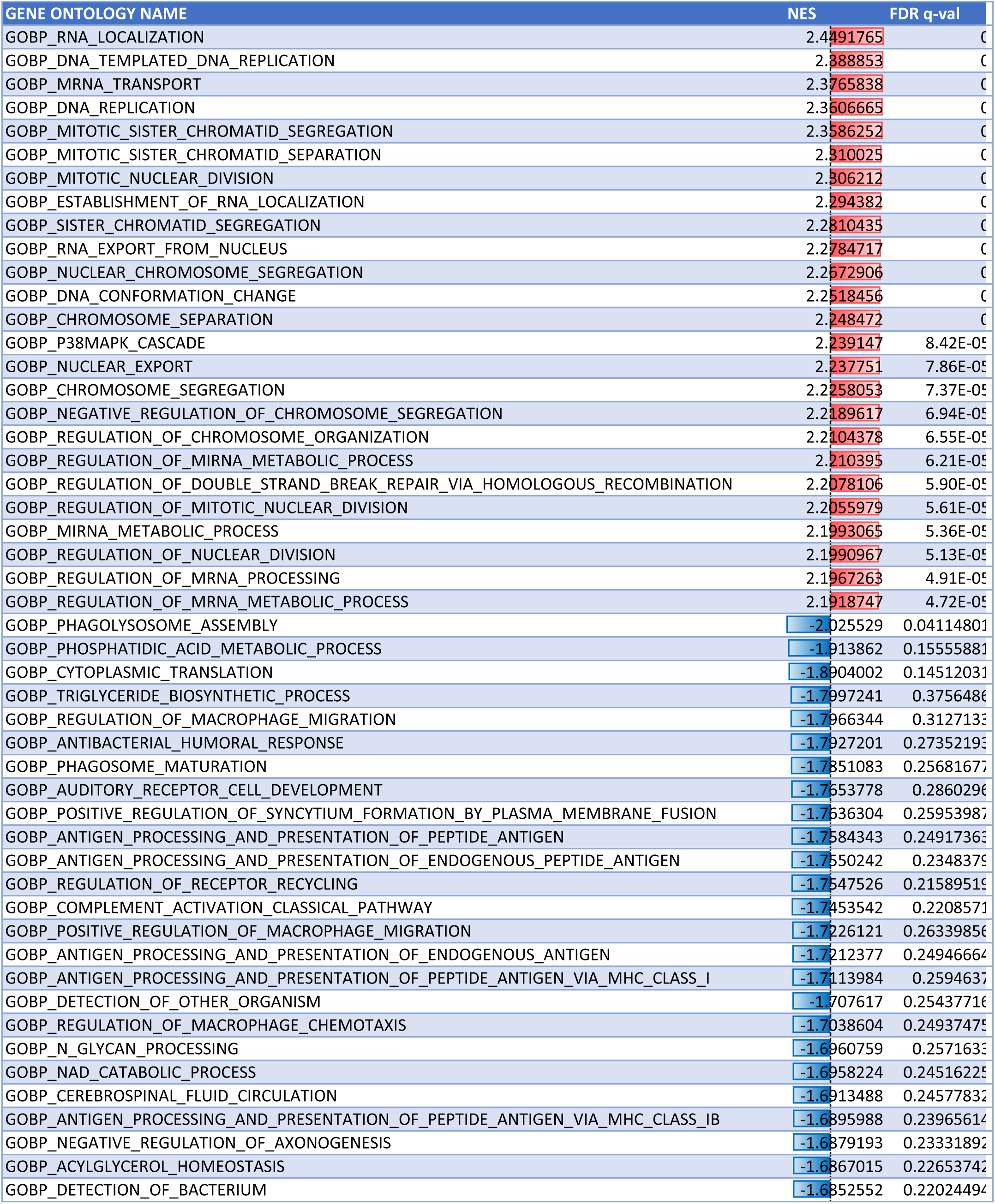
Gene set enrichment analysis from bulk RNA-seq analysis. Related to Figure 4. Gene set enrichment analysis from bulk RNA-seq of 2µm confined DCIS cells. The top 25 upregulate and downregulated gene ontologies are shown.

## METHODS

### KEY RESOURCES TABLE

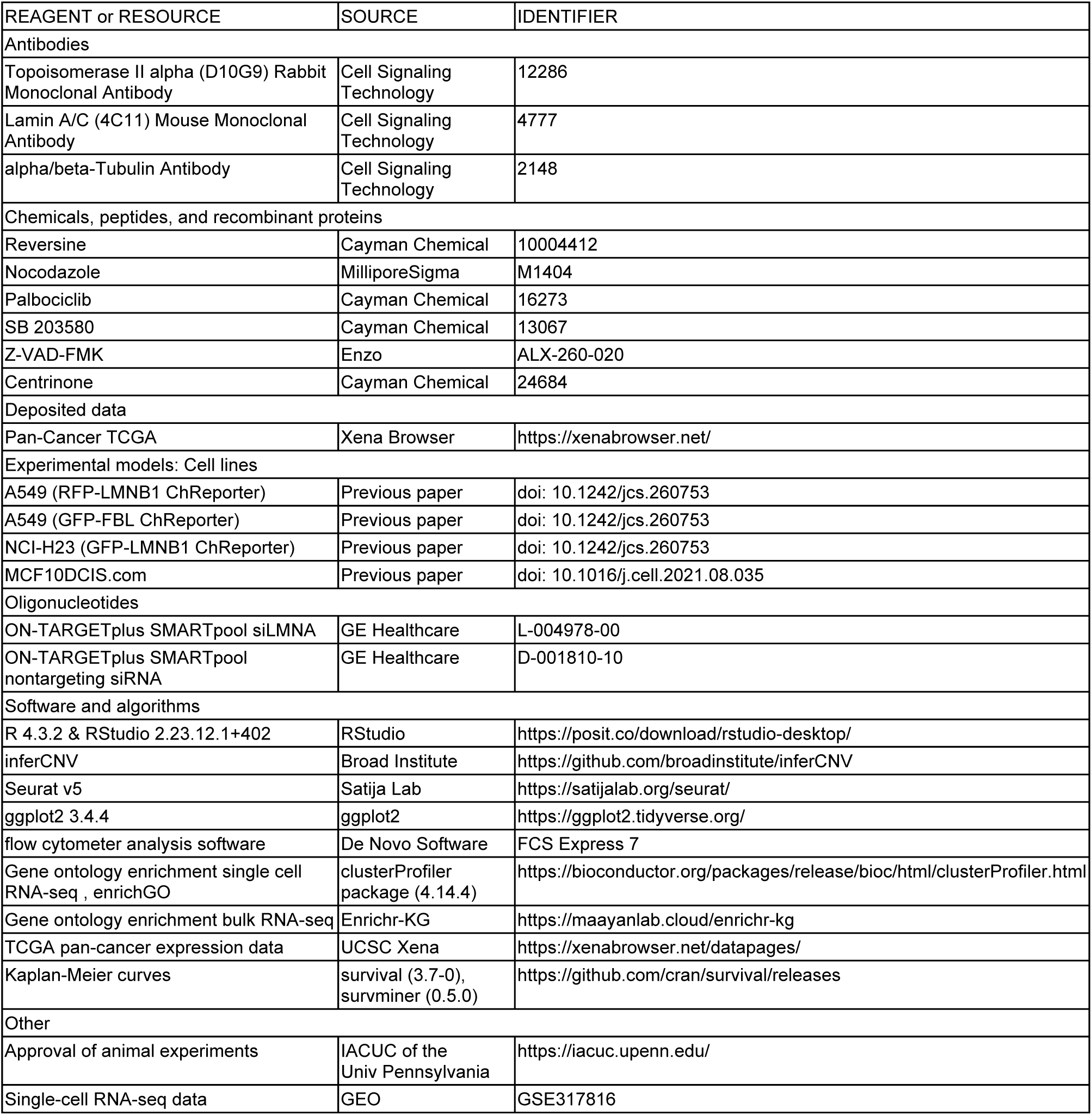

| REAGENT or RESOURCE | SOURCE | IDENTIFIER |
| --- | --- | --- |
| Antibodies |  |  |
| Topoisomerase II alpha (D10G9) Rabbit Monoclonal Antibody | Cell Signaling Technology | 12286 |
| Lamin A/C (4C11) Mouse Monoclonal Antibody | Cell Signaling Technology | 4777 |
| alpha/beta-Tubulin Antibody | Cell Signaling Technology | 2148 |
| Chemicals, peptides, and recombinant proteins |  |  |
| Reversine | Cayman Chemical | 10004412 |
| Nocodazole | MilliporeSigma | M1404 |
| Palbociclib | Cayman Chemical | 16273 |
| SB 203580 | Cayman Chemical | 13067 |
| Z-VAD-FMK | Enzo | ALX-260-020 |
| Centrinone | Cayman Chemical | 24684 |
| Deposited data |  |  |
| Pan-Cancer TCGA | Xena Browser | https://xenabrowser.net/ |
| Experimental models: Cell lines |  |  |
| A549 (RFP-LMNB1 ChReporter) | Previous paper | doi: 10.1242/jcs.260753 |
| A549 (GFP-FBL ChReporter) | Previous paper | doi: 10.1242/jcs.260753 |
| NCI-H23 (GFP-LMNB1 ChReporter) | Previous paper | doi: 10.1242/jcs.260753 |
| MCF10DCIS.com | Previous paper | doi: 10.1016/j.cell.2021.08.035 |
| Oligonucleotides |  |  |
| ON-TARGETplus SMARTpool siLMNA | GE Healthcare | L-004978-00 |
| ON-TARGETplus SMARTpool nontargeting siRNA | GE Healthcare | D-001810-10 |
| Software and algorithms |  |  |
| R 4.3.2 & RStudio 2.23.12.1+402 | RStudio | https://posit.co/download/rstudio-desktop/ |
| inferCNV | Broad Institute | https://github.com/broadinstitute/inferCNV |
| Seurat v5 | Satija Lab | https://satijalab.org/seurat/ |
| ggplot2 3.4.4 | ggplot2 | https://ggplot2.tidyverse.org/ |
| flow cytometer analysis software | De Novo Software | FCS Express 7 |
| Gene ontology enrichment single cell RNA-seq , enrichGO | clusterProfiler package (4.14.4) | https://bioconductor.org/packages/release/bioc/html/clusterProfiler.html |
| Gene ontology enrichment bulk RNA-seq | Enrichr-KG | https://maayanlab.cloud/enrichr-kg |
| TCGA pan-cancer expression data | UCSC Xena | https://xenabrowser.net/datapages/ |
| Kaplan-Meier curves | survival (3.7-0), survminer (0.5.0) | https://github.com/cran/survival/releases |
| Other |  |  |
| Approval of animal experiments | IACUC of the Univ Pennsylvania | https://iacuc.upenn.edu/ |
| Single-cell RNA-seq data | GEO | GSE317816 |

### EXPERIMENTAL MODEL AND STUDY PARTICIPANT DETAILS

#### Cell lines and culture conditions

A549 (CCL-185), H23 (NCI-H23), and U2OS (HTB-96) were obtained from the American Type Culture Collection (ATCC) and cultured in Ham’s F12 (Gibco 11765054), RPMI 1640 (Gibco 11875093), and DMEM GlutaMAX supplement (Gibco 10566016), respectively. Generation and validation of ChReporter cell lines were described previously.^19^ MCF10DCIS.com were cultured in Advanced DMEM/F-12 (Gibco 12634010). All media was supplemented with 10% (v/v) fetal bovine serum (FBS, MilliporeSigma F2242) and 100 U/ml penicillin-streptomycin (Gibco 15140122). All cells were passaged every 3-4 days using 0.05% trypsin-EDTA (Gibco 25300054). All cells were incubated at 37°C and 5% CO_2_. Cell lines were authenticated by SNP array and tested negative (1-2 times annually) for mycoplasma contamination.

#### Tumor xenograft studies in mice

A549 xenografts were generated in immunodeficient non-obese diabetic/severe combined (NOD/SCID) mice with null expression of interleukin-2 receptor gamma chain (NSG mice) as procured by the University of Pennsylvania Stem Cell and Xenograft Core. In 8- to 12-week male NSG mice, subcutaneous injections were made with a 100 μl bolus of ∼0.5×106 A549 cancer cells. Palpable tumors after several weeks of growth were harvested for chromatin height measurements: tumors were fixed overnight using 4% paraformaldehyde at 4°C, permeabilized using 0.5% (v/v) Triton-X for 1 h at room temperature, and finally stained with Hoescht 33342 overnight at 4°C. Tumor slices in PBS were imaged using a Leica TCS SP8 system with a 63×/1.4 NA oil-immersion. All images were taken every 0.5 μm along the focus (Z-stack) to cover the entire nuclei, whether interphase or mitotic. All image stacks were 3D-reconstructed using ImageJ, and the thinnest portion of the chromatin identified as the height.

#### Intravital imaging of mitosis in mouse skin

3-week-old male C57BL/6 mice were anaesthetized with IP injection of ketamine and xylazine, the skin around the head regions were shaved, and the mice were placed on a heated stage with their heads and ears supported by a custom-made stage. A glass coverslip is placed against the skin in the junction region between the head and ear. Image stacks of the skin are acquired with a LaVision TriM Scope II (LaVision Biotec) microscope equipped with ChameleonVision II (Coherent) two-photon laser (940nm for GFP, 1040nm for RFP) focused through a water immersion lens (1.0 NA) and scanned over a 0.5mm^2^ field at 600 Hz. Optical sections in 2µm steps image a total depth of 100µm of tissue in 5min intervals. Anesthesia is maintained throughout with isofluorane by nose cone.

We confirm that all mouse experiments conform to the relevant regulatory standards and maintenance/care, institutional permissions and oversight as approved by the IACUC of the University of Pennsylvania. Effects of sex were not studied and a limitation of the study although A549 cells are male derived.

### METHOD DETAILS

#### Confinement

Polydimethylsiloxane (PDMS, RTV615-010) micropillars of the desired height were prepared, and confinement was performed as previously described.^11^ Briefly, plasma-treated 13/25mm glass coverslips were placed on a 10/1 w/w, PDMS/Crosslinker mixture on wafer molds with micropillars of the desired height. The setup was baked at 95°C for 20 minutes, and then the coverslips with the PDMS micropillars were removed from the wafer. Coverslips were passivated with pLL/PEG (SuSoS, PLL(20)-g[3.5]-PEG(2)) in 10mM HEPES buffer at room temperature for 1 hour, and then incubated in medium overnight before confinement. Large PDMS pillars, holding the coverslips with the micropillars, were attached to a modified 6 well plate lid. This confinement lid was then lowered onto the 6 well plate to push the micropillars on top of a droplet of cells.

#### Drug Treatments

The following drug treatments were used: MPS1 inhibitor reversine (A549: 50nM, H23: 50nM, U2OS: 0.5µM) (Cayman Chemical 10004412), Nocodazole (0.1µM or 0.3µM) (MilliporeSigma M1404), CDK4/6 inhibitor Palbociclib (10µM) (Cayman Chemical 16273), p38 inhibitor SB 203580 (10µM) (Cayman Chemical 13067), pan-caspase inhibitor Z-VAD-FMK (20µM) (Enzo ALX-260-020), and centrosome inhibitor centrinone (0.125µM) (Cayman Chemical 24684).

#### FACS and Flow Cytometry

To prepare cells for fluorescence-activated cell sorting (FACS) and flow cytometry, cells were trypsinized, washed, and resuspended in FACS buffer (5% FBS in PBS). For FACS, cells were additionally passed through a cell strainer (Fisher Scientific 08-771-23). FACS was performed on a BD FACSAria Fusion flow cytometer (Benton Dickinson) to enrich for ChReporter-positive cells. To analyze ChReporter loss following experiments, cells were run on a BD LSRII (Benton Dickinson) flow cytometer, and data was analyzed using FCS Express 7 (De Novo Software).

#### Microscopy

The following experiments were performed using an Olympus IX inverted microscope with 20x/0.4 NA or 40x/0.6 NA objectives using an sCMOS camera (Photometrics Prime): micronuclei quantification, mitosis quantification, immunofluorescence protein expression. The following experiments were performed using a Zeiss LSM 900 with Airyscan 2 with 20x/0.8 NA, 40x/1.2 NA water immersion or 63x/1.4 NA oil immersion objectives: live imaging of cells during and after confinement, ChReporter-negative colonies, nuclear rupture quantification, micronuclei quantification, mitosis quantification.

#### Transfection

All siRNAs used in this study were purchased from GE Healthcare (ON-TARGETplus SMARTpool siLMNA L-004978-00; nontargeting siRNA D-001810-10). Following FACS enrichment of ChReporter-positive U2OS cells, an siRNA pool (25nM) with 1µg/ml Lipofectamine 2000 (Thermo Fisher Scientific 11668030) was created following the manufacturer’s protocol, added to cells, and left for a 3-day incubation. Knockdown efficiency was confirmed with immunofluorescence staining.

#### Immunofluorescence staining

Cells were fixed with 4% paraformaldehyde (Thermo Fisher Scientific 28908) for 15 min. Next, they were permeabilized with 0.5% Triton-X (MilliporeSigma 112298) for 15 min. Then, they were blocked with 5% bovine serum albumin (BSA, MilliporeSigma A7906) for 30 min. Following overnight incubation in primary antibody (1:500 dilution in 5% BSA), cells were incubated in secondary antibody (1:500 dilution) for 1h, and then stained with 8µM Hoechst 33342 (Thermo Fisher Scientific 62249) for 15 min. The following primary antibodies were used: TOP2A (Cell Signaling Technology 12286), LMNA/C (Cell Signaling Technology 4777), Tubulin (Cell Signaling Technology 2148). The following secondary antibodies were used: donkey anti-mouse or rabbit 488, 546, 647 (Invitrogen)

#### Single cell RNA-sequencing

RNA libraries were constructed using the Chromium Single Cell Gene Expression kit (v3.1, single index, Catalog no. PN-1000128; PN-1000127; PN-1000213) from 10X Genomics per the manufacturer’s instructions. The libraries were submitted to the University of Pennsylvania’s Next Generation Sequencing Core for sequencing using NovaSeq 6000 (100 cycles) from Illumina. Raw base call (BCL) files were analyzed using CellRanger (version 5.0.1) to generate FASTQ files and the ‘count’ command was used to generate raw count matrices aligned to GRCh38 provided by 10X genomics.

#### Bulk RNA-sequencing

RNA was isolated using the Qiagen RNeasy RNA extraction kit (Catalog no. 74104) following the manufacturer’s protocol. RNA was then submitted to the University of Pennsylvania’s Next Generation Sequencing Core for library preparation and sequencing using 6000 (100 cycles) from Illumina. The reads were aligned to GRCh38.

### QUANTIFICATION AND STATISTICAL ANALYSIS

Please describe here all statistical analysis and software used.

#### Bioinformatics analysis

All analysis was performed using R 4.3.2 & RStudio 2.23.12.1+402. InferCNV (https://github.com/broadinstitute/inferCNV) was used to construct the copy number profiles following the standard pipeline. UMAPs were generated using Seurat v5. Differential expression analysis was performed using FindMarkers function in Seurat. Gene ontology enrichment analysis of scRNA-seq was done using the clusterProfiler package (4.14.4) and the enrichGO function. Gene ontology enrichment for bulk RNA-seq was done using Enrichr-KG (https://maayanlab.cloud/enrichr-kg). TCGA pan-cancer expression data was downloaded from UCSC Xena (https://xenabrowser.net/datapages/). Clustering was performed using kmeans function in the base library. Kaplan-Meier curves were generated using the survival (3.7-0) and survminer (0.5.0) packages.

#### Statistical Analysis and Curve Fitting

Statistical analysis was performed using the base library of R 4.3.2 & RStudio 2023.12.1+402. All plots were generated using the package ggplot2 3.4.4. Significance was generally assessed as p<0.05, with further statistical indications in figure and table legends.

### ADDITIONAL RESOURCES

None.

